# Concomitant infection with *Leishmania donovani* and *Plasmodium berghei alters* clinical and immune responses in BALB/c mice

**DOI:** 10.1101/2021.04.08.438937

**Authors:** Rebeccah. M. Ayako, Joshua. M. Mutiso, John. C. Macharia, David Langoi, Lucy Ochola

## Abstract

Malaria and visceral leishmaniasis coexist in the same geographical regions. However, dual co-infection with parasites causing these diseases and their impact on public health is poorly documented. Interactions between these parasites may play a role in disease outcome. The present study set out to evaluate the clinical and immunological parameters following *Leishmania donovani* and *Plasmodium berghei* co-infection in BALB/c mice. Mice were divided into four groups; *L. donovani-* only, *L. donovani- P. berghei, P. berghei-* only and naïve. Body weight, parasite burden, total IgG, IFN-γ and IL-4 responses were determined. To determine the survival rate, four mice were used from each group. Tissues for histological analysis were taken from spleen, liver and brain. Results indicated significant differences in body weight (P<0.0001), *L. donovani* parasite load (P< 0.0001*), L. donovani* IgG (P< 0.0001), *P. berghei* parasitemia (P= 0.0222), *P. berghei* IgG (P= 0.002), IFN-γ (P<0.0001) and IL-4 (P<0.0001) in dual-infected mice. There was no correlation between *L. donovani* parasite load and IgG responses in single or dual infections, while there was a positive relationship of *P. berghei* parasitemia and IgG responses in the dual infection group only. *Plasmodium berghei* had the highest mortality rate compared to *L. donovani*- only and *L. donovani- P. berghei* infected mice groups. Histological analyses showed enlarged red and white pulps and pathological changes in the spleen, liver and brain tissues which were less pronounced in co-infected group. We conclude that *L. donovani* and *P. berghei* co-infection reduces disease severity and these changes seem to correlate with variation in serum IgG and cytokines (IFN-γ and IL-4). Therefore, the study recommends the importance of inclusion of early screening of malaria in Visceral Leishmaniasis patients in regions where malaria is co- endemic.

**Author Summary:** Visceral leishmaniasis and malaria are the principal causes of morbidity and mortality affiliated with parasitic diseases universally warranting the necessity to investigate the control and immunology of the infections. Notwithstanding the probable incidences of leishmaniasis- malaria infections in endemic regions are not readily eminent to the clinicians if an individual is co-infected and almost frequently, such patients develop a fever and are customarily treated against malaria and hence the need to study disease progression and outcome during a co-infection. Furthermore, it is unclear if this co-infection could impede the clinical symptoms of the separate diseases and thus the necessity to demonstrate disease outcome in experimentally co-infected murine models. This present study was crucial to find out whether this mode of co-infection alters disease progression and enhanced severity leading to high morbidity and mortality. This current research was an imperative step in using murine as a model in the study of disease outcome and immunopathogenesis of visceral leishmaniasis and malaria co-infection thus establishing the feasibility of co-infecting the BALB/c mice with *Leishmania donovani* and *Plasmodium berghei*.

## Introduction

Visceral leishmaniasis (VL) commonly known as kala-azar is a systemic disease transmitted by phlebotomine female sandflies (1). It is affecting millions of people globally and comes second after malaria in terms of casualties caused by parasitic diseases (2). It is caused by *Leishmania infantum* in the new world and *L. donovani* in the old world and is almost always fatal if left untreated (3). East Africa (Kenya, Ethiopia, Somalia, Sudan and South Sudan), South-East Asia Region (Pakistan, Bangladesh and India), South America (Brazil, Colombia and Peru) and Mediterranean region (Saudi Arabia, Afghanistan and Algeria) account for over 90% of the total cases globally (4).

Malaria is a parasitic disease infecting humans and is transmitted by female mosquitoes of the genus *Anopheles* (5). *Plasmodium* species are the cause of human malaria deaths which were 450,000 with 218 million cases in 2017 and 228 million in 2018 (6). Africa still holds a high proportion of the global malaria burden recording 96% malaria cases and 94% of malaria deaths in global statistics (7).

Belachew *et al*., (2018) showed that in mice infected with *L. donovani*, the balance between Th_1_ and Th_2_ influences disease resistance and susceptibility. Further studies have demonstrated that cytokines induced by Th_1_ responses are linked to efficacious leishmaniasis control while Th_2_ induced cytokines are associated with disease severity (8). Interferon γ (IFN- γ), during malaria and leishmanial infections, is confederated with protective immunity (9).

Co-infections of malaria and VL are the leading cause of increased mortality and morbidity associated with parasitic infections globally [4, 8]. However, regions where co- morbidity occurs, the impact in public health and the outcome of either diseases is poorly documented. Both infections present similar clinical signs and symptoms and many challenges remain to be overcome among patients living in co-endemic areas. Moreover, in human studies due to co-infection, the immune responses and the role they play in reduction of disease severity and protection is not clearly elucidated. High morbidity and mortality associated with the two diseases warranted us to study the disease outcome during a co-infection. Furthermore, it is unclear whether a co-infection could interfere with the clinical outcomes of the diseases. To better understand the clinical and immunological manifestation of a co-infection, we designed the present study to further elucidate the parameters and the effect they have in disease outcome in BALB/c mouse model.

## Methods

### Experimental design

The present study involved a total of 82 BALB/c mice of either sex aged 6-8 weeks which were acquired from the Rodent facility, Animal Science Department at the Institute of Primate Research (IPR [www.primateresearch.org]), Karen, Nairobi, Kenya and housed in the same premises throughout the experimental period. The mice were housed in cages, the room was at an ambient temperature of 22°C and relative humidity of 50-70%. They were fed on mice pellets (Unga Farm Care, Nairobi, Kenya) and water was provided *ad libitum*. The BALB/c mice were divided into 4 groups as follows: *L. donovani- only* infected mice (n= 25); *L. donovani- P. berghei* co-infected mice (n=22) (which were co-infected with *P. berghei* 60 days post-*L. donovani* infection); *P. berghei*-only infected mice (n= 25) and naïve (n=10) (which were given PBS only) (Fig 1). All infections were done intraperitoneally with 1 × 10^6^ *P. berghei* and/ or *L. donovani* parasites suspended in 100μl of PBS. *Leishmania donovani* inoculation was termed day -60 while *P. berghei* inoculation was day 0. Body weight measurements were done every day throughout the co- infection/ experimental period. Following co-infection, 3 mice were sacrificed on day 0, 4 and daily from the 6^th^ day till the end of the experimental period from both co-infected and single infected mice groups to obtain blood and spleen samples. Whole blood was used to prepare serum for IgG, IFN**-γ** and IL-4 quantification while tail prick blood samples which were done daily were used for parasitemia determination in *P. berghei* groups. Spleen samples were obtained in all *L. donovani* groups to prepare splenic impression smears for evaluation of *L. donovani* parasite load. In the survivorship experiment, BALB/c mice were euthanized using 3L/min Carbon IV Oxide (CO_2_) for 2 minutes when they appeared moribund and date recorded. Spleen, liver and brain tissues were harvested on the 9^th^ day from both experimental and control groups for histological analysis. The experimental protocols and procedures were approved by the Institutional Science and Ethics Committee (ISERC) of the Institute of Primate Research (study ISERC/10/2016) and carried out according to its guidelines for Animal care and Handling.

**Fig 1:**
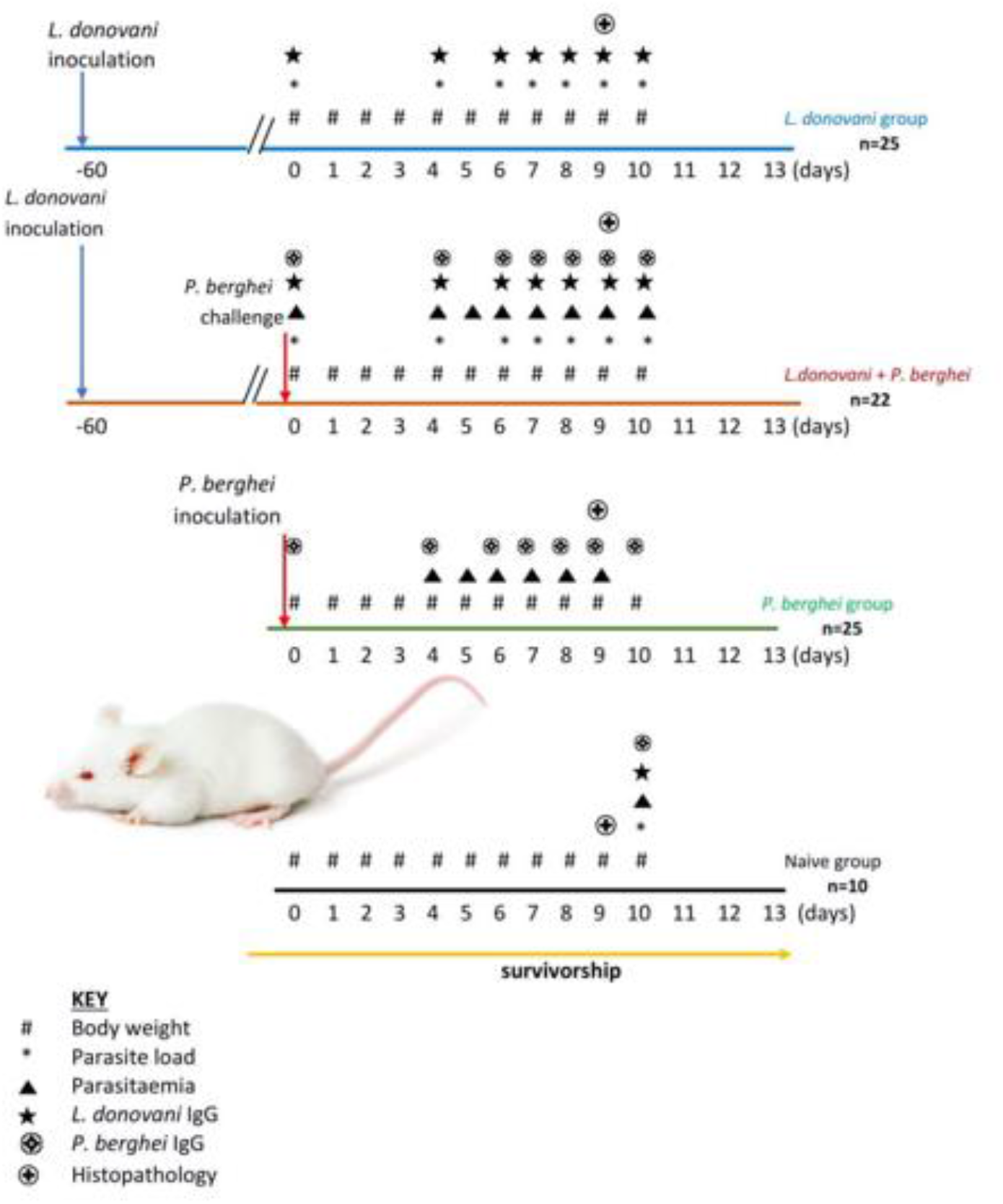
Schematic diagram of the study’s experimental design. Groups of mice were infected with either *L. donovani, L. donovani-P. berghei* or *P. berghei*. A naïve mice group was also included in the study.

### Experimental parasites

*Leishmania donovani* strain NLB-065 was obtained from hamster’s spleen by aspirate and cultured in complete Schneider’s Insect media (Sigma-Aldrich Laborchemikalien GmbH, Seelze, Germany). Promastigotes at the stationary phase were harvested by centrifugation at 2,500 rpm at 4°C for 15 minutes as described (11). The pellet was later washed in sterile Phosphate Buffered Saline (PBS) by centrifugation as before. These parasites were then used to infect mice and for Soluble *Leishmania* antigen (SLA) preparation.

Cryopreserved stocks of *Plasmodium berghei ANKA* (wild type, supplied by Malaria Research and Reference Reagent Resource Center program (MR4)) were retrieved from liquid nitrogen and revived by thawing at 37°C in a water bath followed by three washes in PBS at 2000 rpm for 10 minutes. The parasites were used to infect the BALB/c mice and for malaria crude antigen preparation used in antibody quantification using ELISA.

### Preparation of soluble *Leishmania* antigen (SLA)

The pellet of harvested promastigotes was enumerated and 1 x 10^8^ promastigotes suspended in 2ml PBS, followed by 3 cycles of freeze-thawing in liquid nitrogen and 4 sonication cycles of 18 AMP on ice for 20 seconds each. The parasite suspension was later centrifuged at 10,000g for 30 minutes at 4°C. The supernatant was collected and protein concentration quantified using a Bio-Rad protein assay kit (BioRad Laboratories, USA) and later stored at -80°C until use.

### Preparation of *Plasmodium berghei* antigen

*Plasmodium berghei* antigen was prepared from parasitized Red Blood Cells (pRBCs) obtained from BALB/c mice (12). Four ml of heparinized blood was washed 3 times with PBS at 2000 rpm for 6 minutes. One ml of 0.1 % saponin in PBS was added to the pellet to lyse pRBCs and incubated at room temperature for 10 minutes with occasional mixing. Eight ml of PBS were added and centrifuged at 3000 rpm for 30 minutes at 4°C. The supernatant and RBC ghost were removed and the lysate washed 3 more times until it was dark red. The suspension was sonicated 6 times for 10 seconds with a 1-minute interval in ice to lyse parasite cell wall. The lysate was transferred to microcentrifuge tubes and centrifuged at 14,000 rpm for 60 minutes at 4^0^C. The supernatant, which was the antigen, was harvested and filter sterilized. Protein concentration was determined using the Bio-Rad kit (BioRad Laboratories, USA).

### Body weight measurement

The body weights were measured for both the experimental and control groups daily using a digital laboratory weighing balance (Amazon, USA) and results recorded in grams.

### Parasite burden determination

For *P. berghei* infected BALB/c mice parasite density was determined by examining a Giemsa-stained thin smear prepared from a tail blood prick, daily beginning on day 4 post-*P. berghei* infection. Parasitaemia was determined at 100X oil immersion under a microscope. Uninfected and infected red blood cells were identified and counted from different fields of view. About 500 erythrocytes were counted per slide and parasitemia was calculated as described (13). For *L. donovani* amastigotes, splenic impression smears from sacrificed mice were prepared. The fixing of slides and staining procedures was similar to the preparation of *P. berghei* for parasitemia above. Parasites were counted per 500 splenic cell nuclei at 100X oil immersion under a microscope (14).

### Quantification of IgG using ELISA

Anti-parasite antibodies were performed as described (11). Briefly, polystyrene Micro-ELISA plates (Nunc, Copenhagen, Denmark) were coated overnight with 100 μl per well of antigens at a concentration of 10 μg/ml and 5μg/ml for SLA and malaria crude antigen respectively in PBS pH 9.6. The plates were washed 5 times with 200 μl/well 0.05% Tween_20_ in PBS (washing buffer) to remove excess antigen and later blocked with 200 μl/ well of 1% bovine serum albumin (BSA) in washing buffer and incubated at room temperature for 1 hour. The plates were washed 5 times as above to remove excess blocking buffer and 100 μl of the serum samples were added to the wells and the plate incubated at room temperature for 2 hours. The plates were washed again 5 times with the washing buffer to discard unbound serum and 100 μl of horseradish peroxidase-conjugated goat anti-mouse IgG (Sigma-Aldrich, USA) diluted at 1:2000 in 1% BSA in washing buffer was added to each well followed by 1 hour incubation at room temperature. The plates were washed 5 times as before and 100 μl of Tetramethylbenzidine ((TMB), ThermoFisher, USA) substrate added to each well. Optical densities were read at 630 nm using an ELISA reader (Dynatech Laboratories, Sussex, UK) after 15-minute incubation in the dark.

### Quantification of IFN-γ and IL-4

Polystyrene micro-ELISA plates were coated overnight at 4^0^C with 100μl of mAb AN 18 and mAb 11B11 monoclonal antibodies specific for IFN-γ and IL-4 respectively on separate plates. Bovine serum albumin (0.1%) in PBS/0.05% Tween_20_ buffer (washing buffer) was used to block nonspecific binding sites for 1 hour at 37^0^C. The plates were washed 3 times with washing buffer before the addition of 100 μl of serum samples and mouse cytokine standards and incubated for 2 hours at room temperature. Hundred microlitres of monoclonal cytokines detecting antibody, R4-6A2-biotin and BVD6-24G2 (0.5 µg/ml) were added per well and the plate incubated for 1 hour at room temperature. The plates were washed 5 times as before and 100μl Streptavidin (1:1000) added to each well before incubation at room temperature for 1 hour. Unbound Streptavidin was discarded by washing the plates 5 times and 100μl of TMB peroxidase substrate added to each well. The plates were incubated for 30 minutes in the dark and optical densities read at 630nm in a microplate reader (Dynatech Laboratories, UK).

### Determination of survival rate in mice

Following the challenge of BALB/c mice with *L. donovani* and/ or *P. berghei* in the respective groups, four mice from each group were used for the determination of survivorship (15). A survivorship curve was plotted from the percentage of the number of surviving mice for 15 days.

### Histopathology

Following harvesting of the spleen, liver and brain tissues of one mouse from each group on the 9^th^-day post-infection, the organs were fixed in 10% formalin. They were later processed as described (16). Briefly, the tissues were dehydrated in alcohol at increasing concentrations for 1 hour before being immersed in xylene and embedded in melted paraffin wax for 3 hours. The sections were later cut at 4μm thick using a microtome and mounted on microscope slides where they were stained with hematoxylin-eosin stain (Abcam, USA). After they had air-dried, they were viewed under a light microscope at X400 and X100 magnifications and micrographs of the slides were captured.

### Statistical analysis

Statistical analysis was conducted using GraphPad prism software package version (8.01). Body weight, *P. berghei* parasitemia, *L. donovani* parasite load, their respective IgG, IFN-γ and IL-4 responses were calculated as mean ± standard error of the mean (SEM). Mean body weights were compared using a one-way analysis of variance (ANOVA) and adjusted with a Tukey post hoc test. Intragroup statistical analysis was tested using student t-test while independent t-test was used to analyse intergroup means. For correlation analysis, Spearman’s rank correlation test was used. Kaplan-Meier curve was utilized in comparing the different survivorship rates. A difference was considered significant when the *P-*value was less than 0.05 (P < 0.05).

### Ethics

This study and all experimental protocols were approved by the Institutional Science and Ethics Committee (ISERC) of the IPR, Karen, Nairobi, Kenya (study ISERC/10/2016), whose membership is constituted based on guidelines issued by the World Health Organization for committees that review biomedical research, by the NIH, by PVEN, and by the Helsinki Convention on the Humane Treatment of Animals for Scientific Purposes. The IRC-ACUC is nationally registered by the National Commission for Science, Technology, and Innovation, Kenya.

## Results

### Body weights of BALB/ c mice

Following co-infection, variations in body weight were observed throughout the experimental period. Body weight changes in the co-infected and control groups at the termination date ranged from (30.415-21.22g) which was not significantly different (F= 0.8427; P= 0.5402). The mean body weight of *L. donovani-P. berghei* co-infected mice group ranged from 32.4g on day 0 to 28g on the 10^th^ day which showed a significant decrease (P=0.0001). There was a slight drop in the mean body weight of *L. donovani-*only infected mice group from 32.2g on day 0 to 30.4g at the end of the experimental period (P=0.0001). *Plasmodium berghei-*only infected mice group had a slight decrease in body weight from 22.55g on day 0 to 21.58g on the 9^th^ day (P=0.0001). There was however an increase in mean body weight in the naïve mice group from 22.72g to 23.27g (Fig 2) at the end of the experimental period which was (P=0.710).

**Fig 2:**
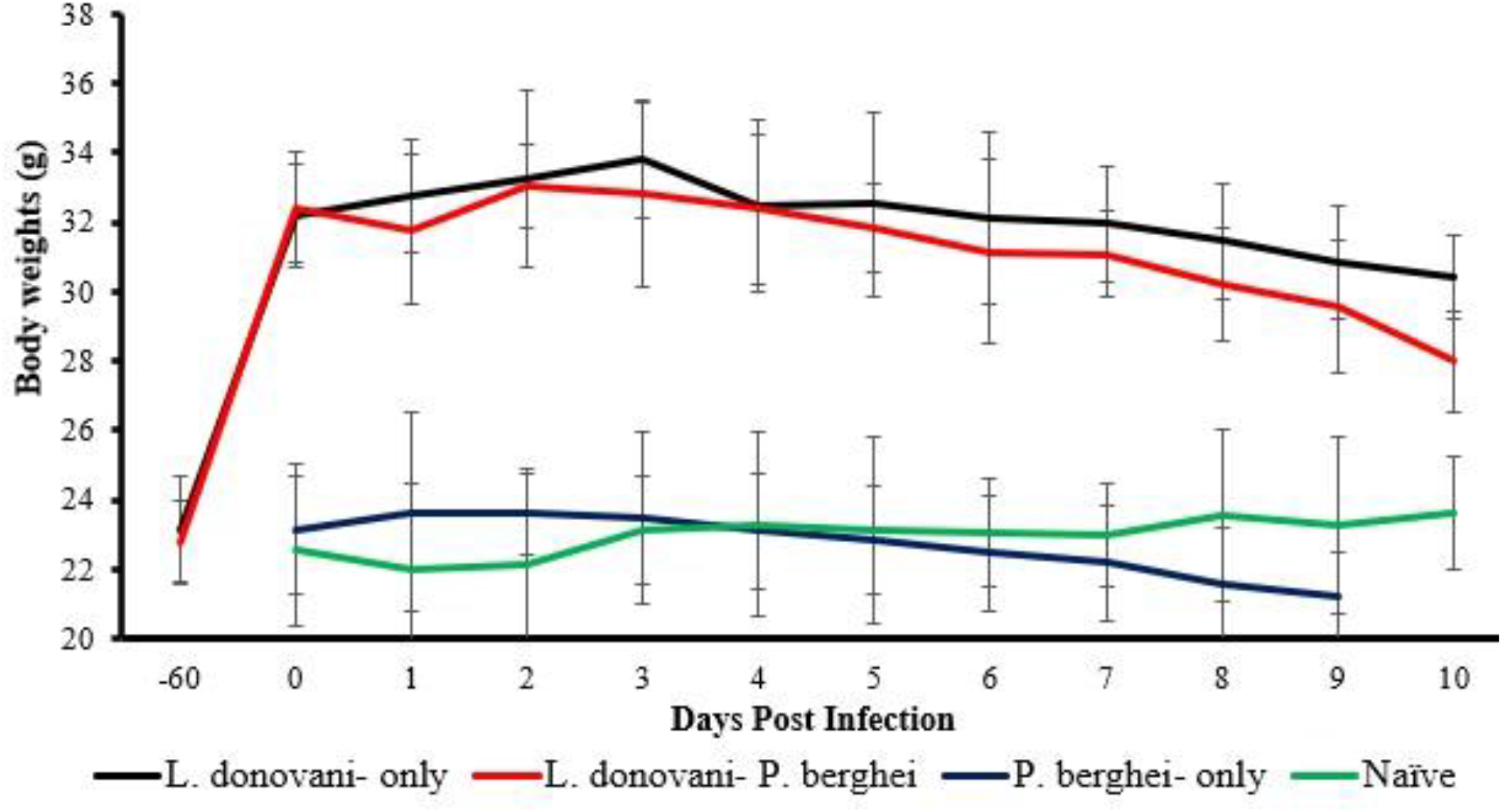
Mean (± SD) body weights of BALB/c mice over 10 days of the experimental period. Groups of mice were infected with either *L. donovani, L. donovani-P. berghei* or *P. berghei* and their body weights taken each day post co-infection for 10 days. A naïve mice group was also included in the study. The mean body weight of *L. donovani-P. berghei* co-infected mice group showed a significant decrease and a slight drop in the mean body weight of *L. donovani-*only infected mice group was observed. *Plasmodium berghei-*only infected mice group had a slight and the naïve control had an increase in mean body weight up to the termination of the experiment.

### *Leishmania donovani* and *Plasmodium berghei* parasite burden

Upon evaluating *L. donovani* parasite load and *P. berghei* parasitemia, there was a significant difference in the mean parasite load in *L. donovani-P. berghei* co-infected and *L. donovani-*only infected BALB/c mice groups (P= 0.0021) and mean *P. berghei* parasitemia between *L. donovani-P. berghei* co-infected and *P. berghei-*only infected BALB/c mice (P=0.0099). The mean splenic amastigote load in *L. donovani-P. berghei* co-infected mice group decreased from 9.85% on day 0 of co-infection to 9.56% (Fig 3) on termination date (P<0.0001) while the mean *L. donovani* parasite load in *L. donovani-*only infected mice was 9.85% on day 0 and 17.84% on termination date which was (P<0.0001) hence remarkably different.

**Fig 3:**
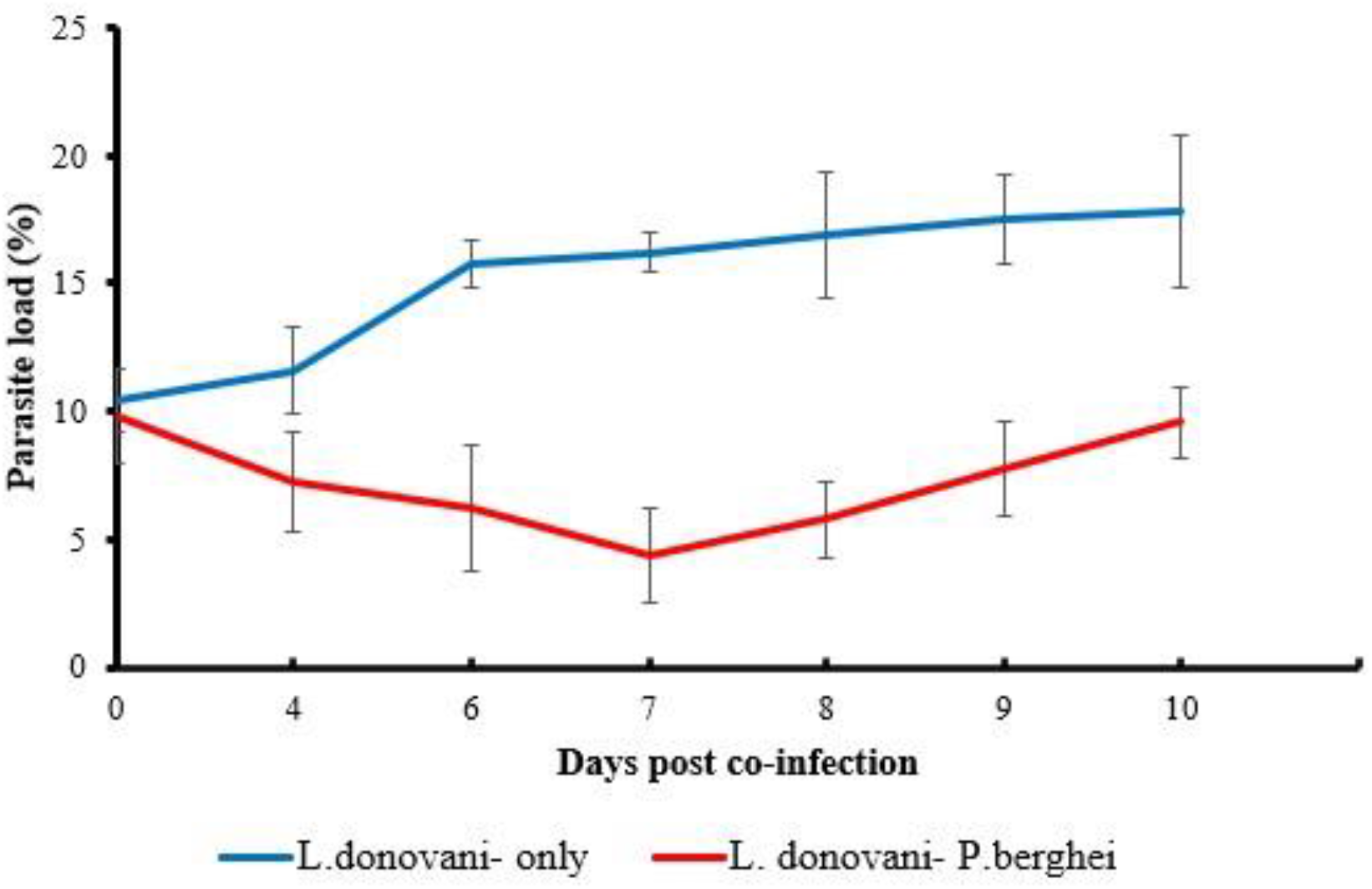
Mean (± SD) parasite load (%) of BALB/c mice over 10 days of the experimental period. Groups of mice were infected with either *L. donovani* or *L. donovani-P. berghei* and their *L. donovani* parasite load taken on day 0, 4 and each day from the 6^th^ day post co-infection. The spleen impression smears were prepared and stained with Giemsa and the parasite load determined by evaluating infected macrophages per 500 splenic nuclei.

The mean *P. berghei* parasitemia in *L. donovani-P. berghei* co-infected mice group was initially 0% on day 4 of co-infection but increased steadily to 14.88% (Fig 4) on day 10 (P= 0.0345). *Plasmodium berghei*-only infected mice had a mean parasitemia of 2.82% on day 4 and on day 9 the parasitemia had markedly increased to 23.45% (P= 0.0222).

**Fig 4:**
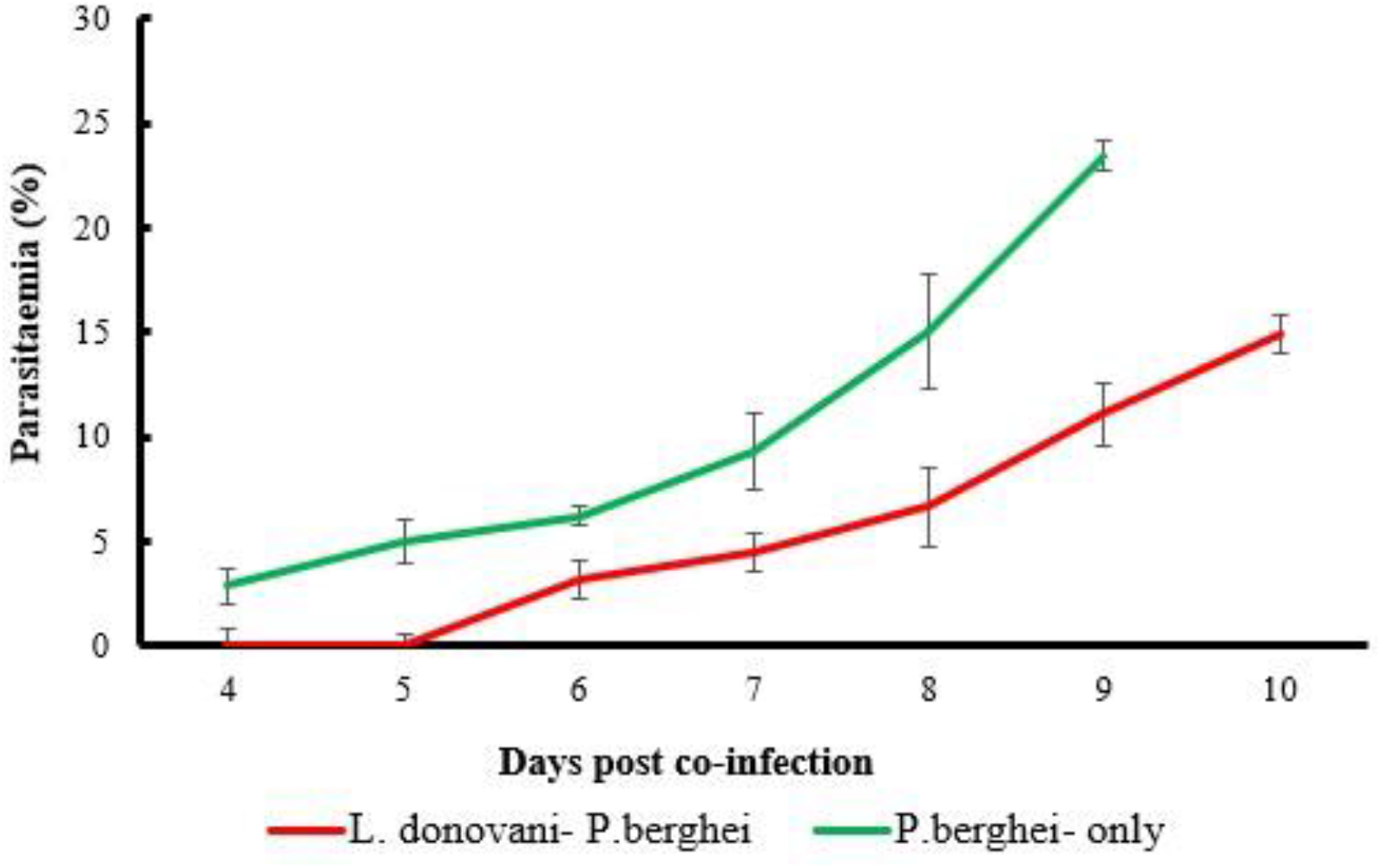
Mean (± SD) parasitemia (%) of BALB/c mice over 10 days of the experimental period. Groups of mice were infected with either *L. donovani-P. berghei* or *P. berghei* and their *P. berghei* parasitemia determined by preparing thin smears of the tail blood on a slide and later staining with Giemsa. The slides were observed using oil immersion at X100 objective lens. Tail blood was taken each day from the 4^th^ day post co-infection.

### *Leishmania donovani* and *Plasmodium berghei* Immunoglobulin gamma (IgG) responses

Following the quantification by ELISA of total IgG specific responses in mice, the mean *Leishmania* specific optical density (OD) compared in *L. donovani-P. berghei* co-infected BALB/c mice were 0.8873 on day 0 and 1.0287 on day 10 (P< 0.0001). The mean OD level in *Leishmania-*only infected mice group ranged from 0.8873 at the beginning of the co-infection period to 1.2597 on day 10 (P< 0.0001) (Fig 5).

**Fig 5:**
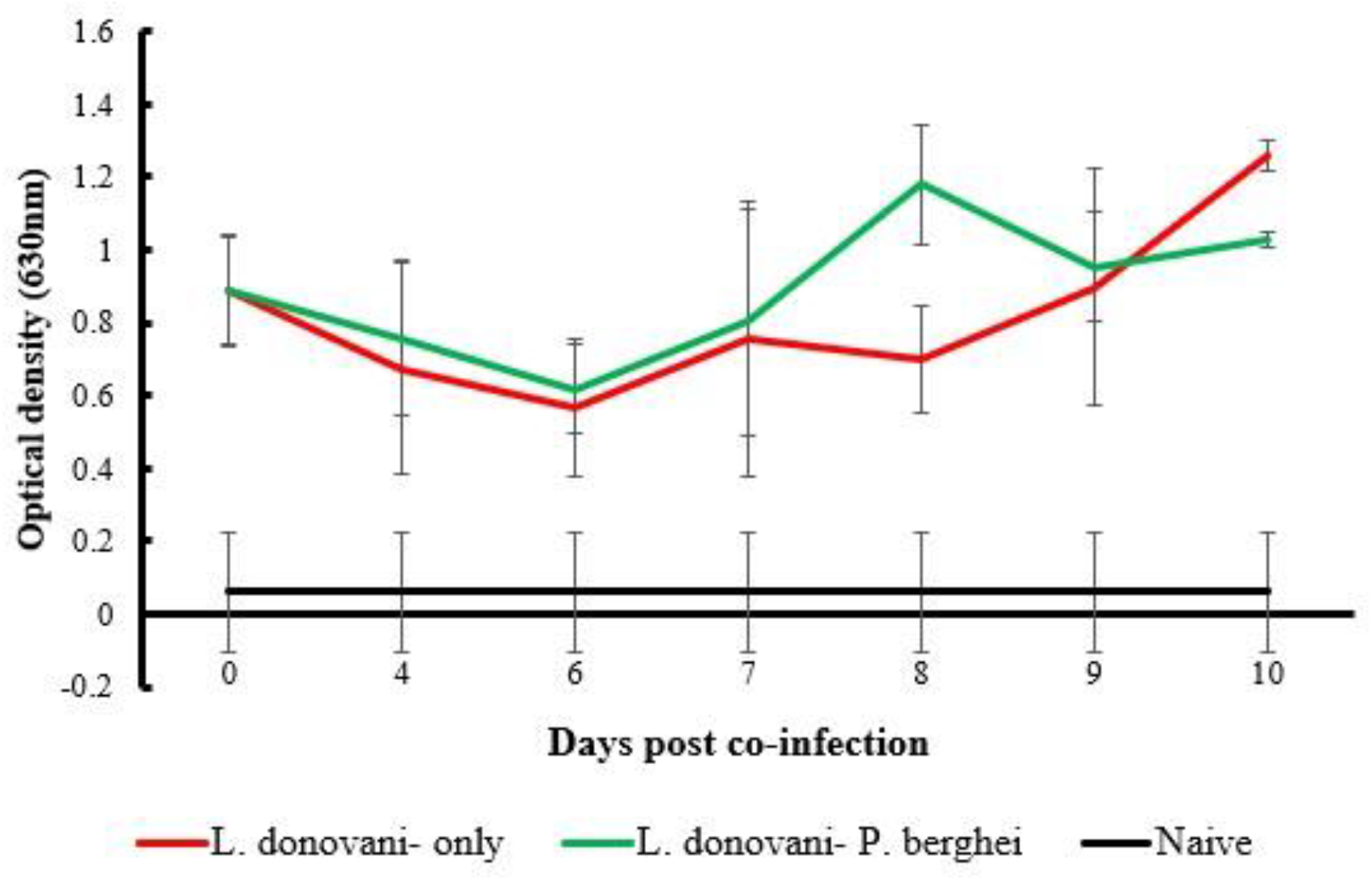
Mean (± SD) *L. donovani* specific IgG optical densities (630nm) of BALB/c mice over 10 days of the experimental period. Groups of mice were infected with either *L. donovani* or *L. donovani-P. berghei* and their IgG optical densities taken on day 0 and each day from the 4^th^ day post co-infection. A naïve mice group was also included in the study. An ELISA plate was coated with Soluble *Leishmania* antigen (SLA) and incubated with serum samples from mice in each group. The optical densities were read at 630nm and the results compared.

The mean *Plasmodium* specific IgG OD levels in *L. donovani-P. berghei* co-infected BALB/c mice group had 0.033 OD at day 0 and 1.285 OD on day 10 (P= 0.002) while mean OD for *P. berghei* only infected mice ranged from OD of 0.032 on day 0 and 0.85 on day 9 (P= 0.0002) (Fig 6).

**Fig 6:**
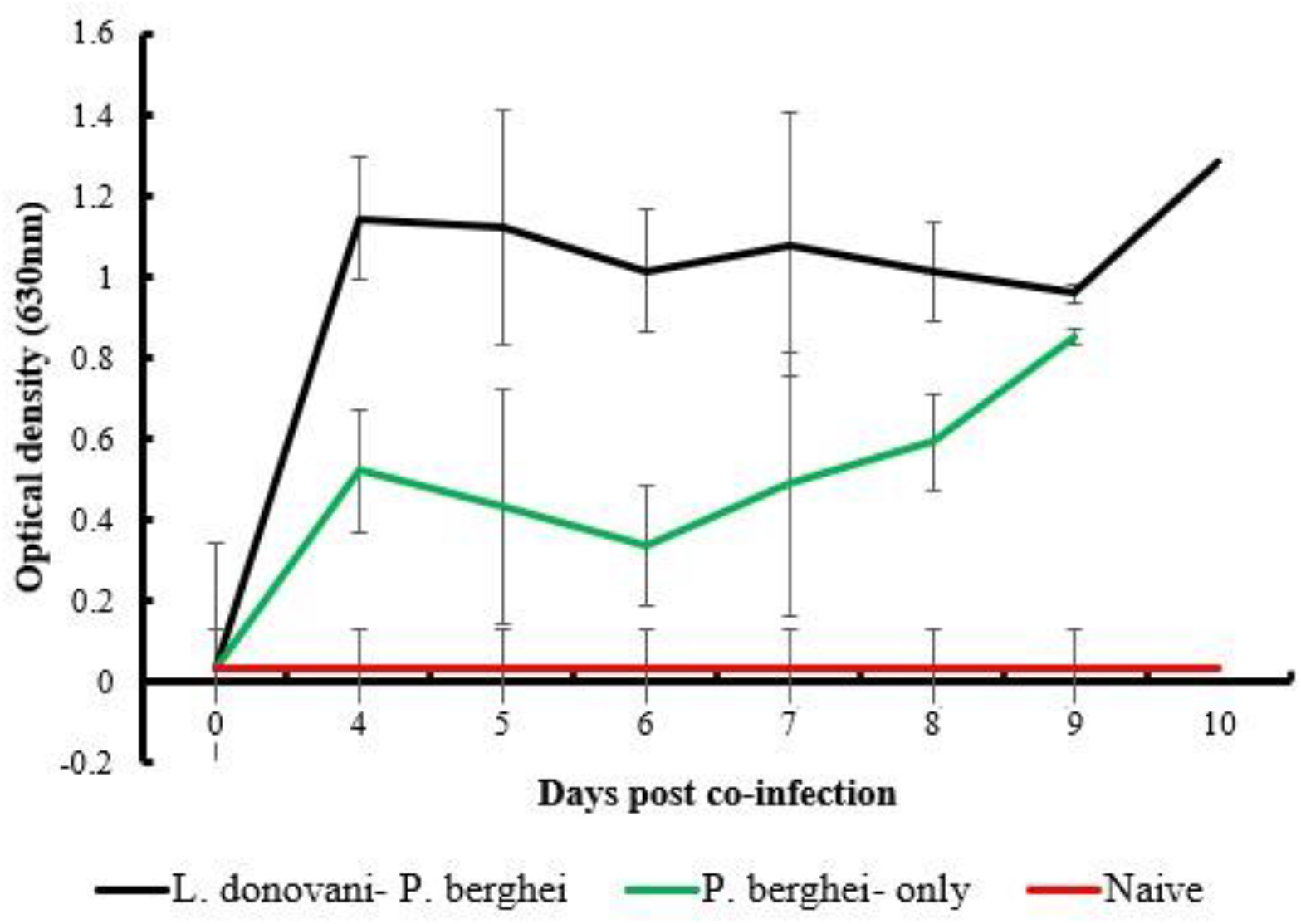
Mean (± SD) *P. berghei* specific IgG optical densities (630nm) of BALB/c mice over 10 days of the experimental period. Groups of mice were infected with either *L. donovani* or *L. donovani-P. berghei* and their IgG optical densities quantified on day 0 and each day from the 4^th^ day post co-infection. ELISA plate was coated with malaria crude antigen. Serum samples were diluted at 1:100 and incubated for 2 hours before the conjugate was added at a concentration of 1:2000 after washing off the unbound antibodies. The substrate was added, the OD read at 630nm using an ELISA reader and mean ODs compared.

### Relationship between antibody responses and parasite burden

The relationship between antibody responses and parasite burden was assessed. Following infection with both *L. donovani* and *P. berghei*, results indicated a decrease in *L. donovani* parasite load coincided with a decrease in *L. donovani* IgG OD values which declined at day 0 post-co-infection to day 6. In general, it was noted that an increase in parasite load resulted in a net increase in OD values as shown from day 7 of co-infection to the end of the experimental period (Fig 7). Similar results were seen in *L. donovani-*only infected mice groups where a decrease in parasite load resulted in a decrease in OD values and vice-versa however these results were not significant ((r= 0.01130 and 0.1816 respectively) P=0.5345).

**Fig 7:**
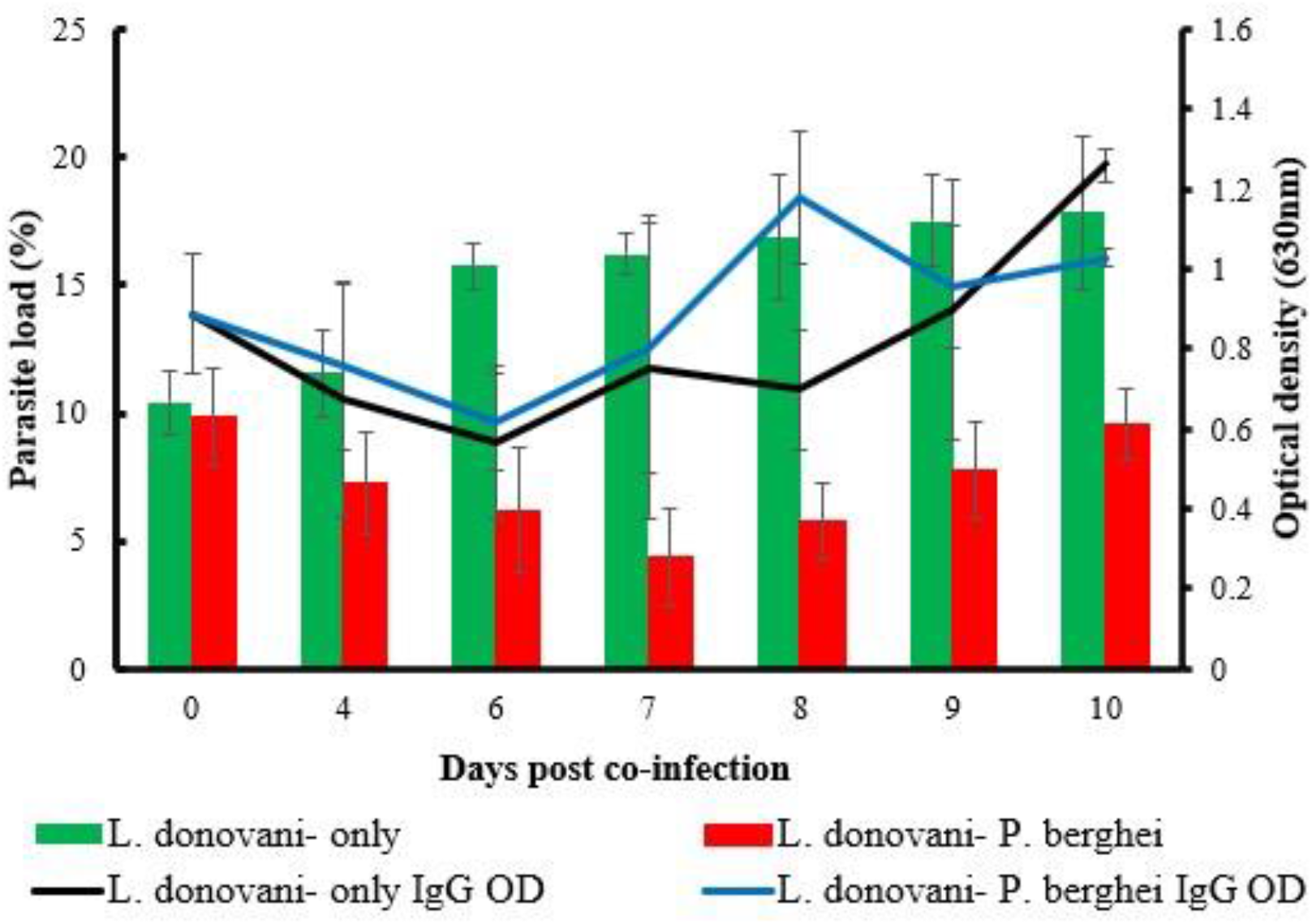
Mean parasite load (%) and *L. donovani* specific IgG optical densities (OD) in *L. donovani-P. berghei* and *P. berghei* infected mice groups. Groups of mice were infected with either *L. donovani* or *L. donovani-P. berghei* and their IgG optical densities with *L. donovani* parasite load established on day 0, 4 and each day from the 6^th^ day post co-infection. An increase in parasite load resulted in a net increase in OD values as shown from day 7 of co-infection when the experiment was terminated.

An increase in parasitemia also coincided with a peak in *P. berghei* IgG antibody responses. High OD values observed in *L. donovani-P. berghei* co-infected mice group resulted in high parasitemia while low IgG responses in *P. berghei* only infected mice group resulted in low parasitemia (Fig 8). A moderate correlation in the mean parasitemia in *L. donovani-P. berghei* and *P. berghei*-only infected mice groups with their respective IgG antibody responses were observed ((r=0.5919 and 0.4833 respectively) P=0.05).

**Fig 8:**
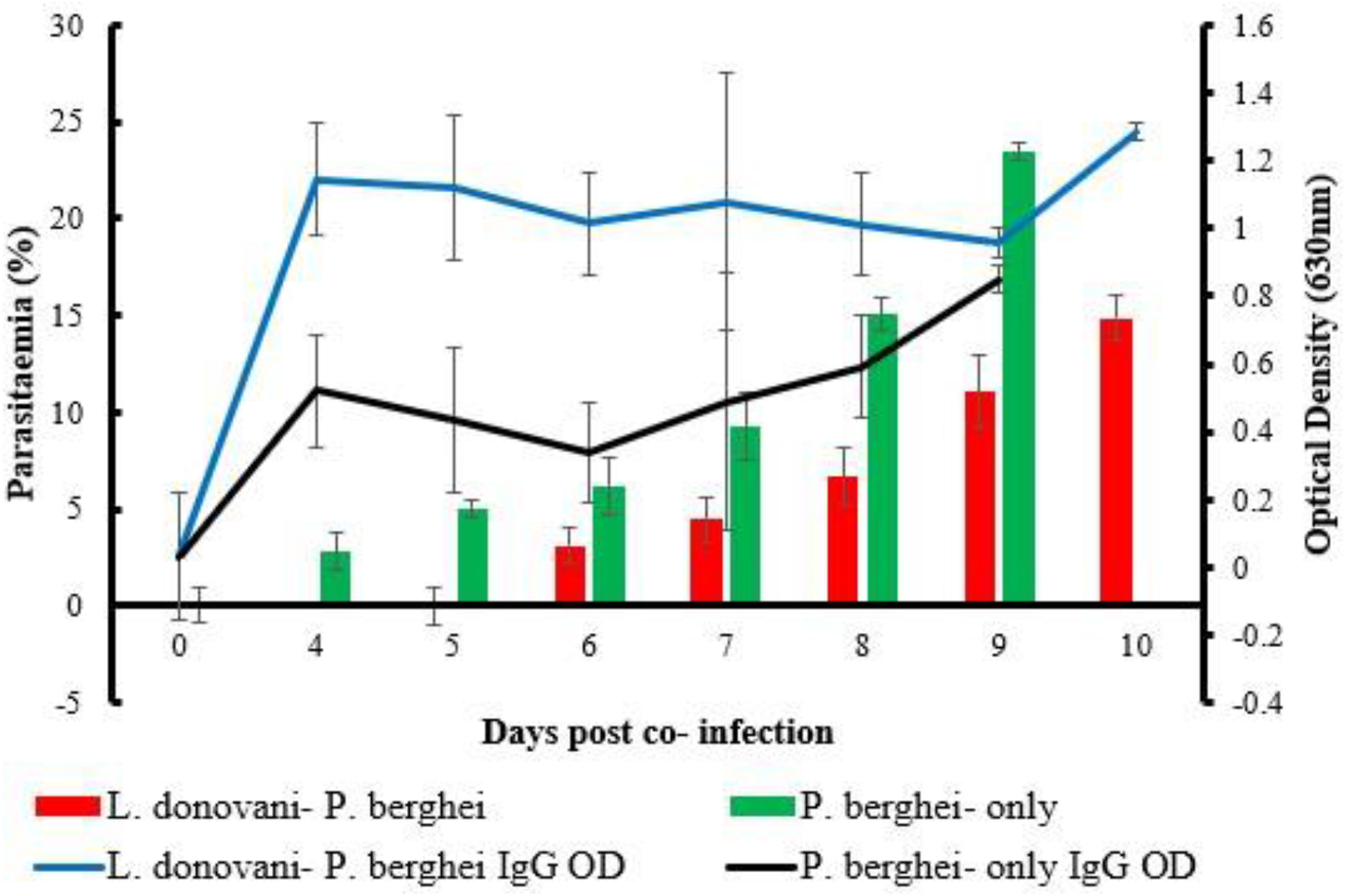
Parasitaemia and optical density (OD) in *L. donovani-P. berghei* and *P. berghei-*only infected mice groups. Groups of mice were infected with either *L. donovani* or *L. donovani-P. berghei* and their IgG optical densities taken on day 0 and each day from the 4^th^ day post co-infection. A moderate correlation in the mean parasitemia in *L. donovani-P. berghei* and *P. berghei*-only infected mice groups with their respective IgG antibody responses were observed.

### Dynamics of (IFN-γ) and IL-4 responses Interferon gamma (IFN-γ) responses

Interferon gamma (IFN-γ) responses of *L. donovani* and *P. berghei* were quantified by measuring ODs by ELISA. We ascertained a decrease in circulating IFN-γ responses in *L. donovani-P. berghei* on day 4 post co-infection before increasing steadily to 0.8653 OD on day 8 (Fig 9). Mean OD started to decrease significantly to 0.4229 on day 10 of co-infection (P= 0.0012). There was a steady decrease in IFN-γ responses in *L. donovani* only (P= 0.0002) while *P. berghei* only infected mice registered a peak in mean IFN-γ levels on day 6 of co-infection after which it began to notably decrease to 0.5264 OD (P= 0.0034). Results also indicated a notable difference in circulating IFN-γ responses within the groups during the co-infection period (P<0.0001).

**Fig 9:**
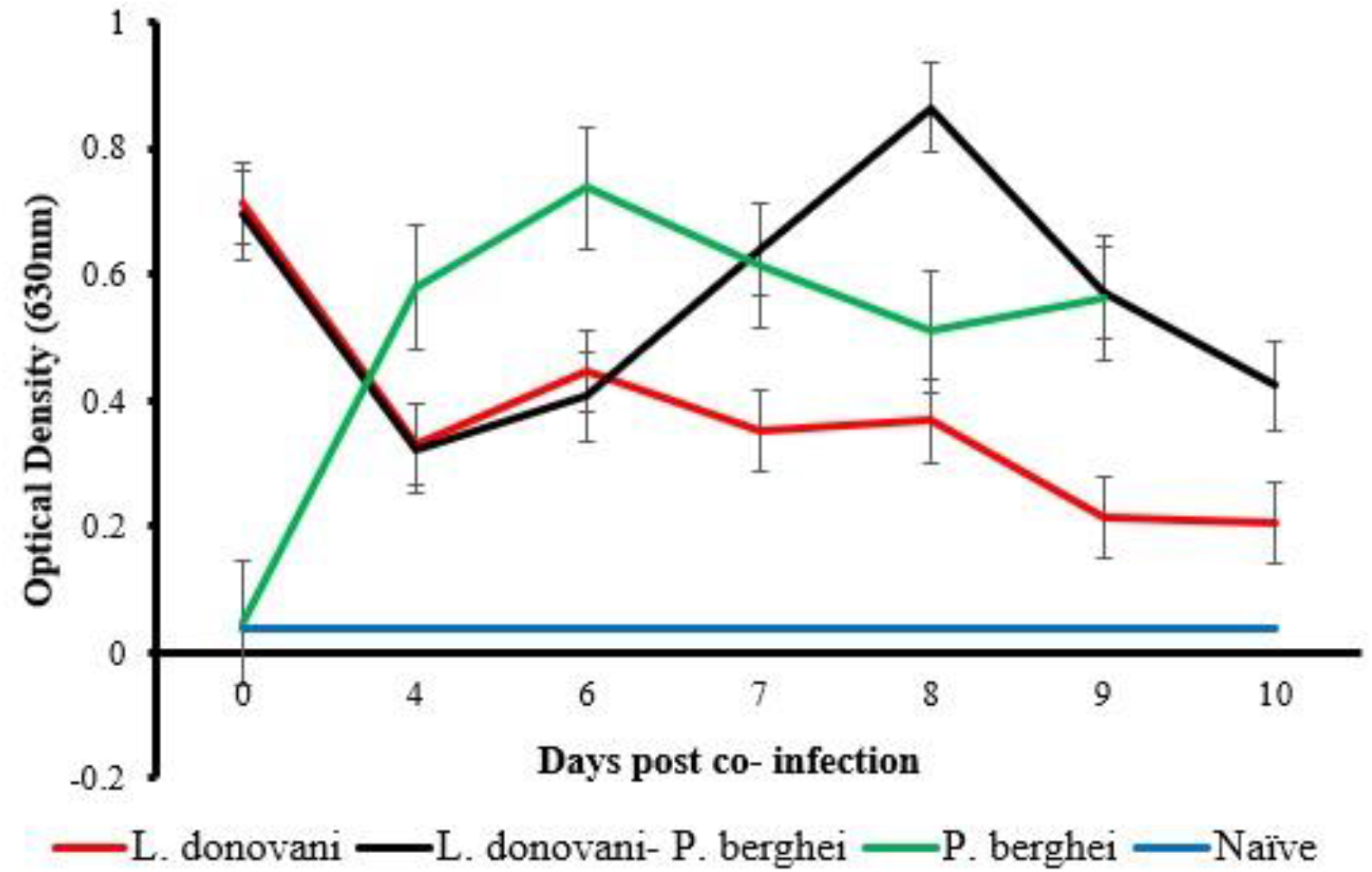
Mean (± SD) IFN-γ optical densities (630nm) of BALB/c mice over 10 days of the experimental period. Groups of mice were infected with either *L. donovani* or *L. donovani-P. berghei* and their IFN-γ optical densities quantified by ELISA reader on day 0 and each day from the 4^th^ day post co-infection. There was a steady decrease in IFN-γ responses in *L. donovani* only while *P. berghei* only infected mice registered a peak in mean IFN-γ levels on day 6 post co-infection.

### Interleukin (IL-4) responses

When we assessed IL-4 responses and it was noted there was a decrease in IL-4 levels from day 0 to day 6 post co-infection. The mean OD levels increased significantly to a high OD of 0.6914 (Fig 10) at the end of the experimental period (P= 0.0001). We also noted a steady significant increase in circulating IL-4 levels in *L. donovani* only (P= 0.0013) and *P. berghei* only (P= 0.0079) infected mice during the experimental period. Data analysis revealed that IL-4 levels were significantly different between the experimental and control groups during the experimental period (P<0.0001).

**Fig 10:**
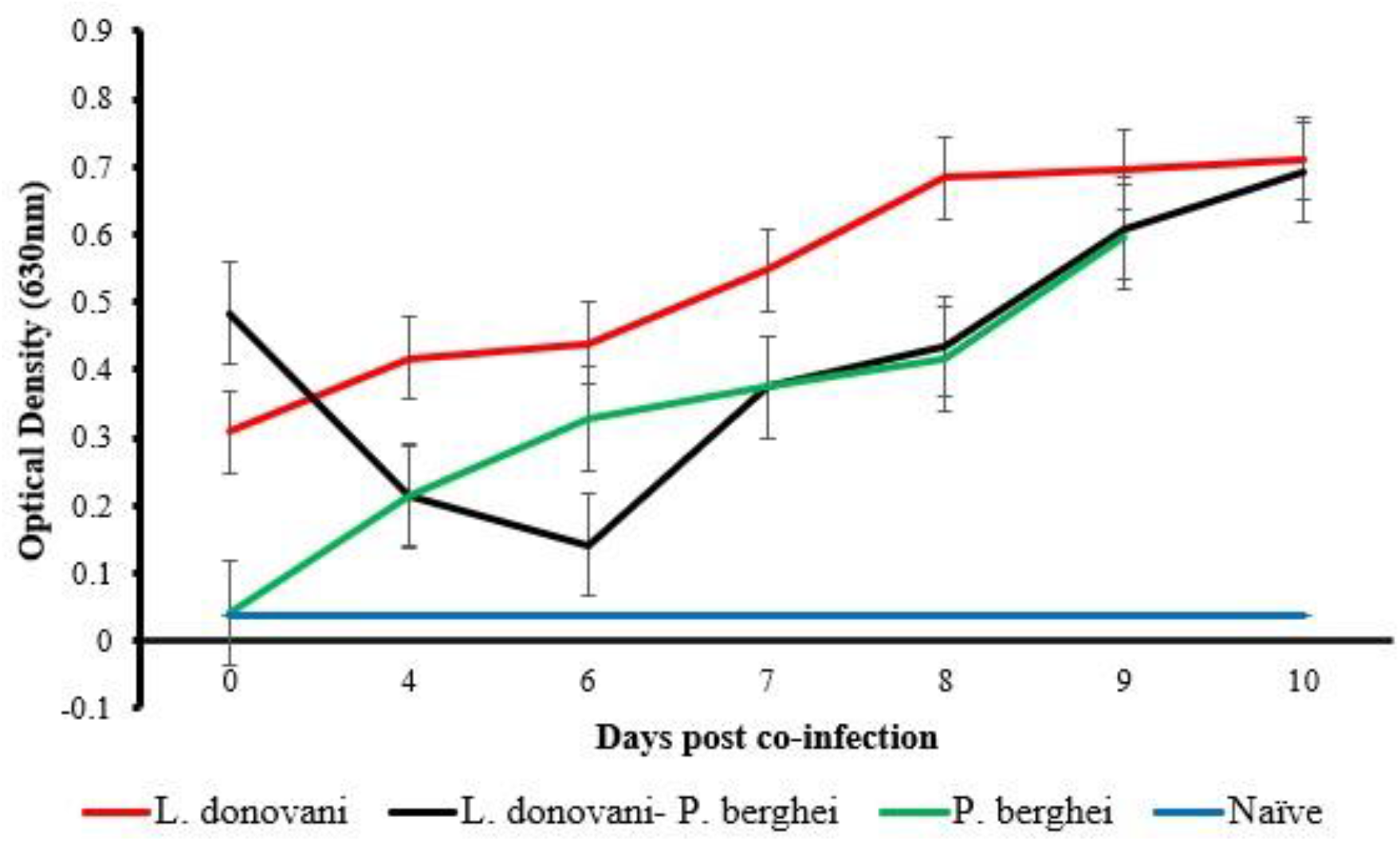
Mean (± SD) IL-4 optical densities (630nm) of BALB/c mice over 10 days of the experimental period. Groups of mice were infected with either *L. donovani* or *L. donovani-P. berghei* and their IgG optical densities taken on day 0 and each day from the 4^th^ day post co-infection. The mean OD levels and IL-4 levels increased significantly in *L. donovani* and *P. berghei* only infected mice during the experimental period.

### Survivorship

The impact of dual and single infections on mice survival was assessed. There were significant differences in the survivorship rate of BALB/c mice in experimental and control groups (P=0.05). The *L. donovani-P. berghei* infected mice survived longer than the *P. berghei* only mice group while *L. donovani* only and naïve mice groups continued to survive up to the 20^th^ day when they were censored. On day 9 post-co-infection, 75% of the *L. donovani-P. berghei* co-infected mice had survived and each mouse was sacrificed on the 10^th^, 12^th^ and 13^th^ day. None of the *L. donovani* only infected BALB/c mice succumbed to infection within the experimental period while 50% of *P. berghei* only infected BALB/c mice were sacrificed on the 6^th^ day and the remaining two were sacrificed on the 7^th^ and 9^th^ day (Fig 14).

### Histopathology

Histological analyses to assess the structural changes of the spleen, liver and brain tissues from all the groups were undertaken. Naïve tissue sections of each organ were also included. The micrographs were viewed at X100 and X400 under a light microscope. Hematoxylin and Eosin-stained micrograph of the spleen showed an enlargement of red and white pulp elements accompanied by a loss in the typical structure of the germinal center with even distribution of granulomas and hyaline deposits in *L. donovani-*only mice which were more evident than *L. donovani-P. berghei* co-infected BALB/c mice. Massive sequestration of parasitized erythrocytes and accumulation of hemozoin was more pronounced in *P. berghei-*only infected mice as compared to *L. donovani-P. berghei* co-infected mice group (Fig 11).

**Fig 11:**
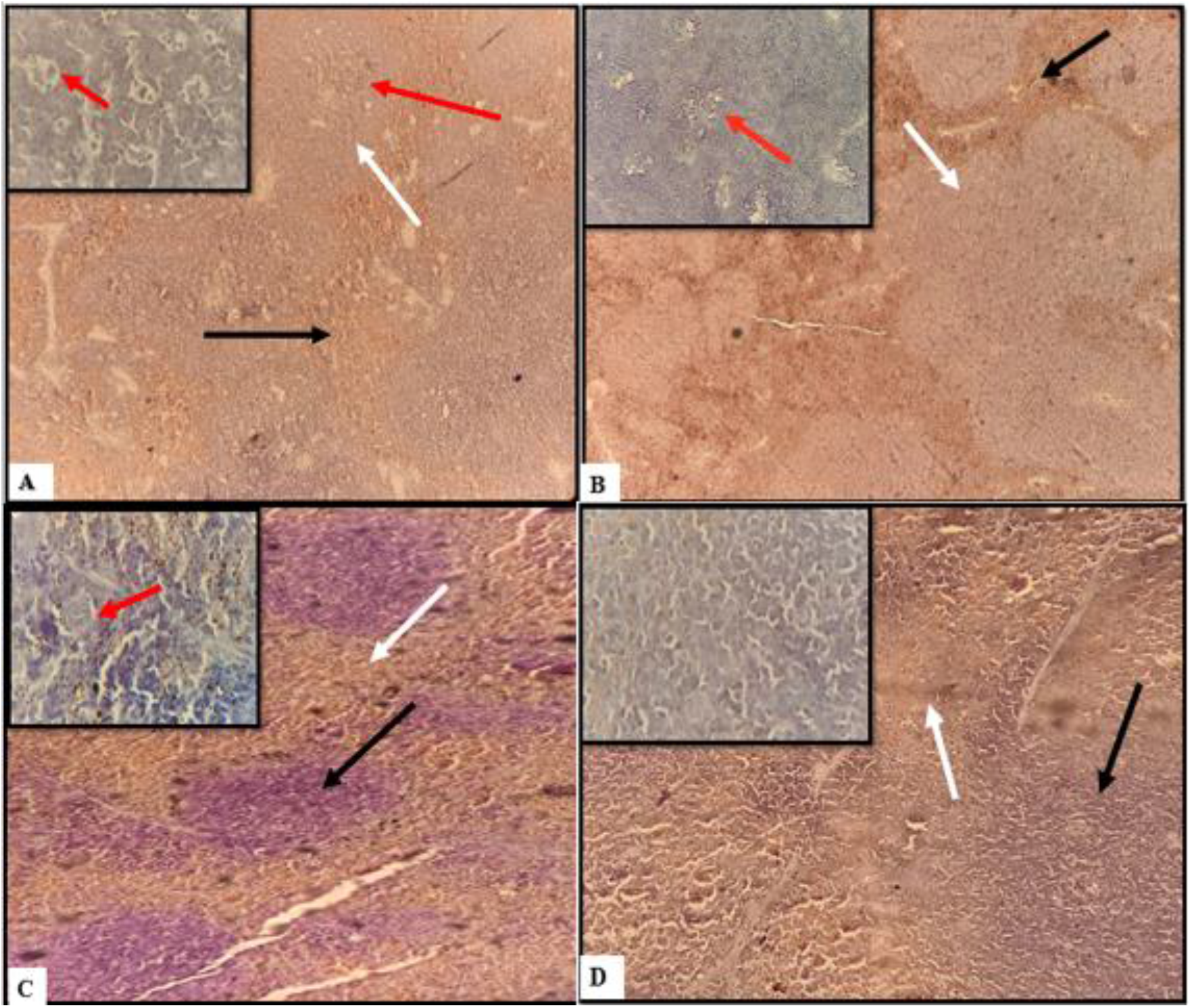
Hematoxylin and Eosin (H&E) stained of spleen section showing structural changes in the white (black arrow) and red pulps (white arrow) with disorganization of granulomas (red arrows). A-*L. donovani-P. berghei* co-infected group; B-*L. donovani* only infected mice group; C-*P. berghei* only and D-naïve group). Viewed under a light microscope at X100 with smaller boxes showing a closer view of granulomas formation (X400 magnification).

H&E-stained liver section revealed lymphocytosis with hyaline deposits and granulomas in *Leishmania only* infected BALB/c mice. Presence of granulomas, necrosis as a result of hepatocyte injury and congestion of hepatic portal vessels was observed in *L. donovani-P. berghei* infected mouse. Multifocal lymphocytes and congestion of blood vessels was evident in *P. berghei* only infected mouse. No histopathological changes were present in the naïve control mice (Fig 12).

**Fig 12:**
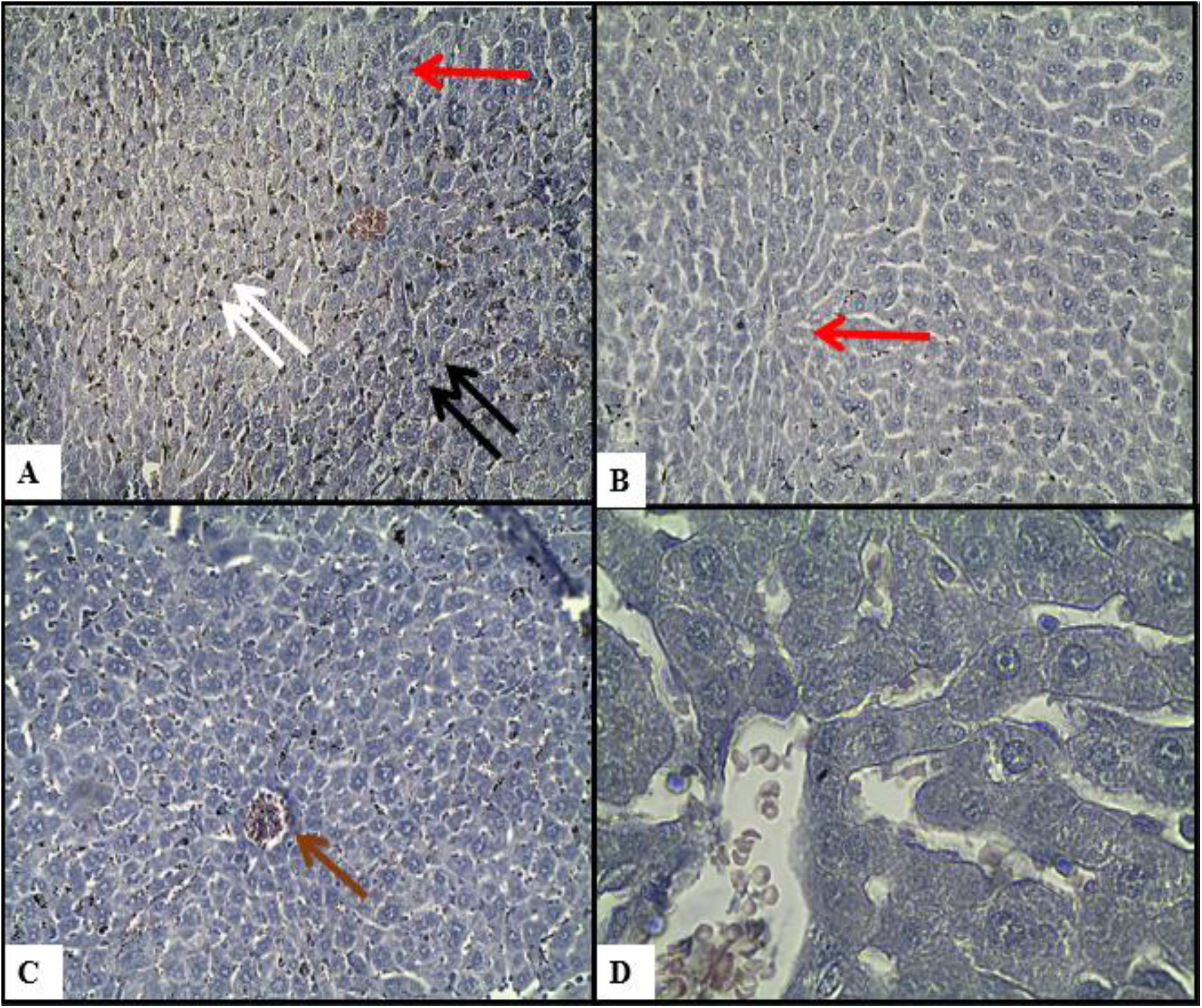
Hematoxylin and Eosin (H&E) stained liver section showing granulomas (red arrows) with congestion of blood vessels (brown arrows) necrosis (black arrows) and hyaline deposits (white arrows). A group of BALB/c mice infected with *L. donovani* and or *P. berghei* parasites before histological analysis of the spleen was carried out at the end of the experimental period. A-*L. donovani-P. berghei* co-infected group; B-*L. donovani* only infected mice group; C-*P. berghei* only and D-naïve group). Viewed under a light microscope at X100 magnification.

H & E-stained sections of the brain were assessed and *P. berghei* only infected mice group showed microhemorrhages and massive leukocyte sequestration were observed while few leukocytes in *L. donovani-P. berghei* co-infected mice were seen. No morphological changes were observed in *Leishmania*-only and naïve control mice (Fig 13).

**Fig 13:**
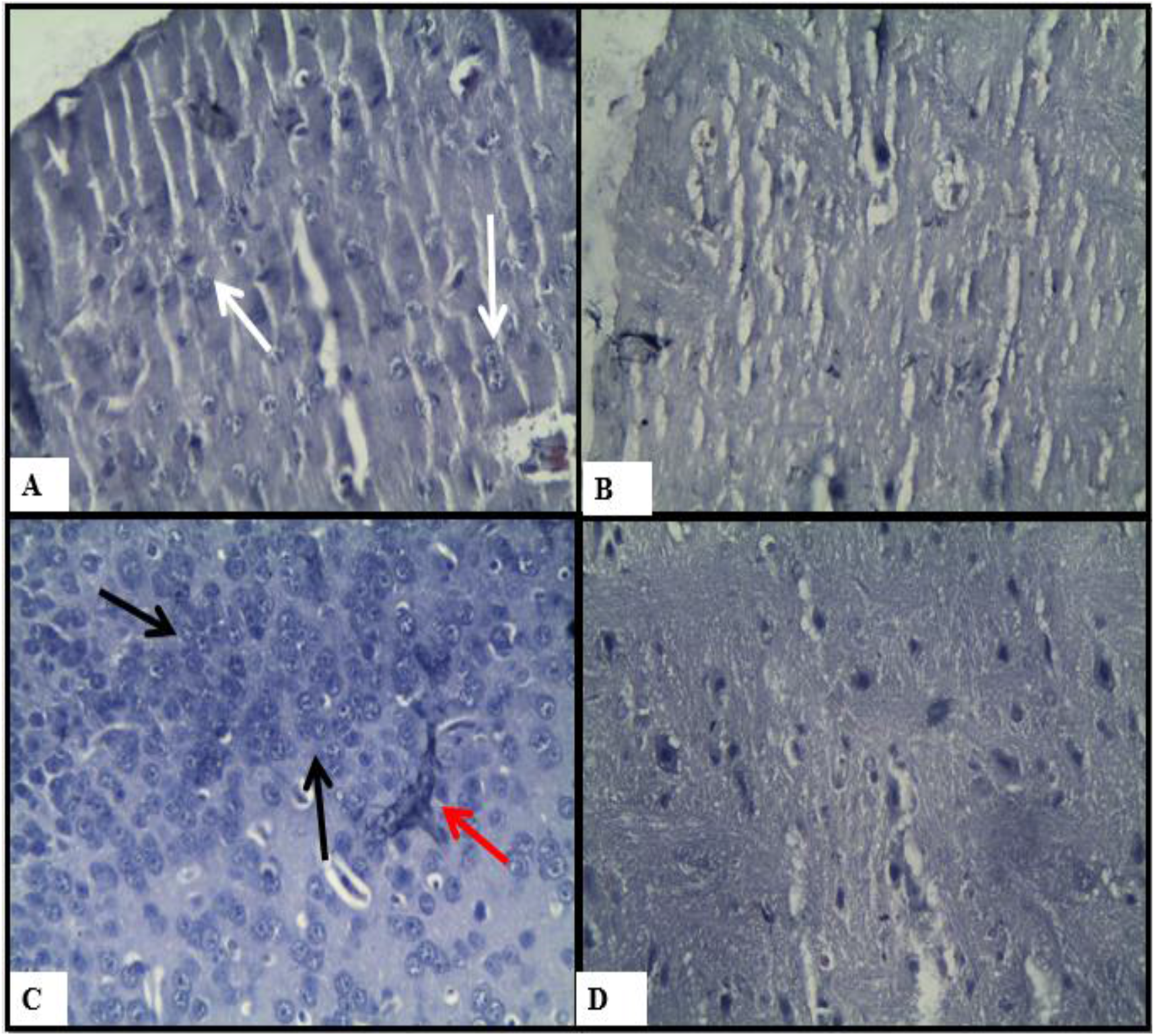
Hematoxylin and Eosin (H&E) staining of brain section showing microhemorrhages (red arrows), massive sequestration of leukocytes (black arrows) and fewer leukocytes (white arrows). (A-*L. donovani-P. berghei* co-infected group; B-*L. donovani* only infected mice group; C-*P. berghei* only and D-naïve group (X400 magnification).

**Fig 14:**
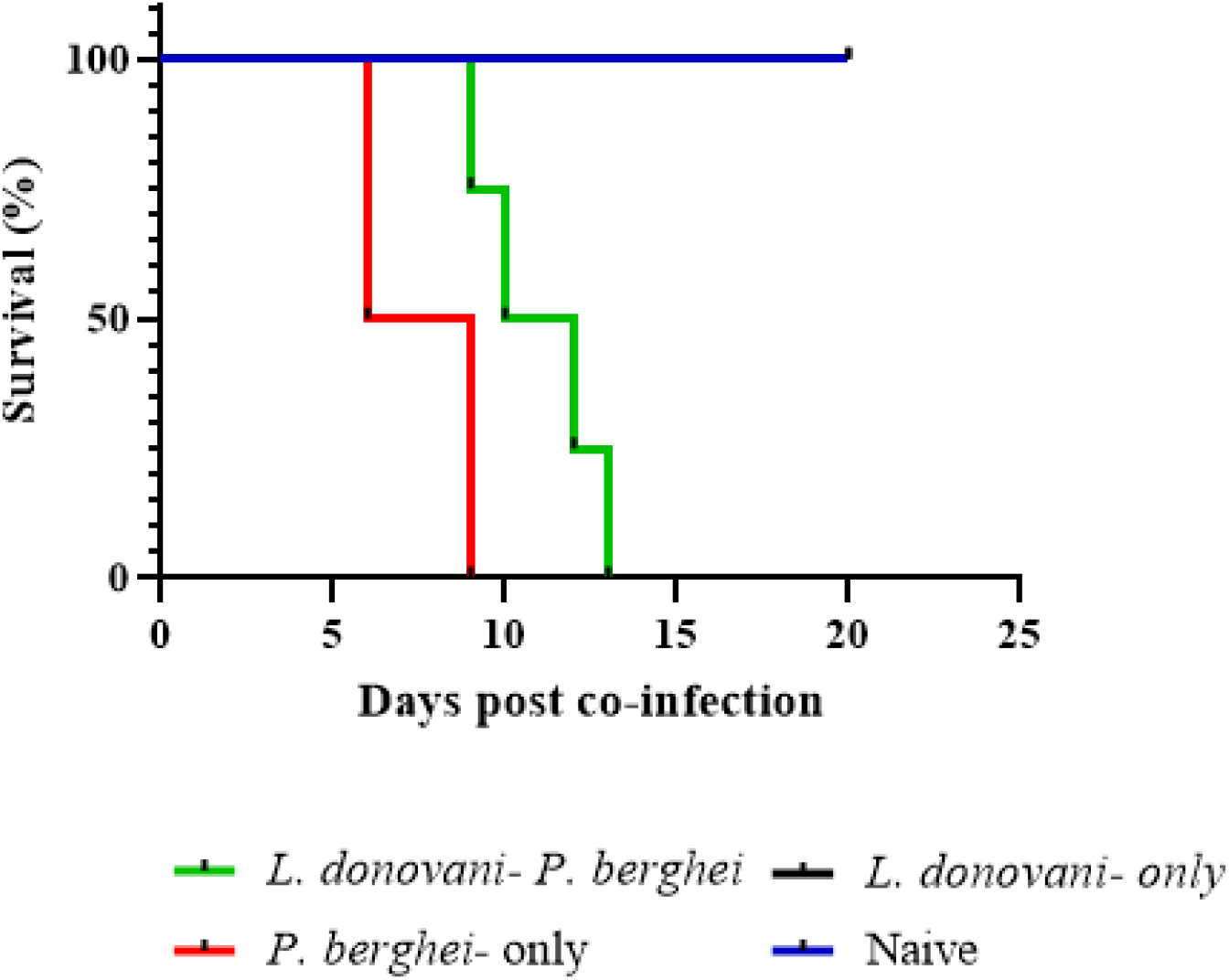
Survivorship curve of BALB/c mice. Groups of mice were infected with either *L. donovani, L. donovani-P. berghei* or *P. berghei* and their survival rate monitored each day post co-infection. *Leishmania donovani-*only and naïve mice groups had a region of overlap. The *L. donovani-P. berghei* infected mice survived longer than the *P. berghei* only mice group while *L. donovani* only and naïve mice groups continued to survive up to the 20^th^ day when they were censored.

## Discussion

The present study was an attempt to evaluate clinical and immunological parameters of *Leishmania donovani* and *Plasmodium berghei* co-infection aimed at establishing the disease outcome in BALB/c mouse model.

There was an overall reduction in body weight in all the infected mice as compared to the non-infected naïve controls. In separate studies, [16, 17] associated diminution in body weight to the loss of appetite, reduced metabolism, altered gut function, hypoglycemia and most importantly massive destruction of erythrocytes. In addition, *P. berghei* infected rats (19) and *L* .*donovani* infected mice (20) studies reported high anemia which was due to the destruction of erythrocytes and splenocytes respectively.

In our study, we established that a delay in the onset and low levels of parasitemia observed in co-infected mice may suggest that pre-infection with *L. donovani* suppresses *P. berghei* infection. While a rapid increase in *P. berghei* parasite density observed during co-infection indicates that increased severity of *L. donovani* predisposes the host to more severe infection. These findings are however not in agreement with (21) where the opposite was the case in that *L. amazonensis* and *P. yoelii* co-infection where parasitemia levels increased on day 5 post-co-infection while *L. braziliensis* and *P. yoelii* co-infection resulted in a decrease and eventual clearance of parasitemia. A decline in *L. donovani* parasite load during the early phase of co-infection in *L. donovani-P. berghei* co-infected mice group implies that superimposing *P. berghei* in *L. donovani* infection may contribute to a reduction in parasite load. In a study to correlate parasitic load with IL-4 responses in patients with cutaneous leishmaniasis, it was noted that parasite burden may play a vital role in effective immune response during pathogenesis (22).

The current study found out that an increase in *L. donovani* parasite load resulted in high IgG antibodies. A study to quantify IgG responses in cutaneous leishmaniasis (CL) patients (23) reported high IgG antibody levels but this was found not to play any role on immunity. In the current study, reduction in IgG responses in co-infected mice which we observed may signify a lower parasite burden as compared to a single infected mice group. IgG plays a substantial role in mediating inhibition of pre-erythrocytic infection in malaria (24). As a result of this, it may explain the elevated IgG antibodies in mice infected with *P. berghei-*only as compared to *L. donovani-P. berghei* co-infected mice group. Previous research by (25) has indicated high antibody titre correlates to high parasitemia levels which is the hallmark in protection against *P. berghei*.

It is documented that erythrocytic stage of malaria infection triggers potent IFN-γ responses in both human (26) and murine studies which explains an increase in IFN-γ responses in the co-infected group during the early phase of co-infection. During this period, we observed low *L. donovani* parasite load and *P. berghei* parasitemia levels suggesting some level of immune protection as a result of superimposed *P. berghei* on *L. donovani* infection. Previous studies revealled that pro-inflammatory cytokines such as IFN-γ, TNF-α and IL-12 secreted by Th1 subset play a significant role in effective control of *L. donovani* infection (27). As malaria infection advanced in the co-infected group, we noticed an increase in anti-inflammatory cytokine (IL-4), a crucial regulatory cytokine secreted in mice during pathology for protection during acute phase of malaria (28) accompanied with a decrease in pro-inflammatory cytokine (IFN-γ). An increase in secreted IL-4 correlated with a decrease in IFN-γ levels and subsequently serum IgG levels. *Leishmania donovani* parasite load and *P. berghei* parasitemia levels increased with an increase in serum IL-4 levels while IFN-γ reduced towards the end of the experimental period suggesting that Th2 induced cytokines are linked to disease severity. From this study, we demonstrated that the balance between Th1 and Th2 determines disease resistance and exacerbation during *L. donovani* and *P. berghei* co-infection.

In areas of co-endemicity, the population are continuously exposed to malaria and leishmaniasis bite hence suffer from possible recurrent infections. This leads to possibility of immunomodulatory effects that may result in interference in the outcome of the two diseases. In the current study, the co-infected group had a higher survival rate compared to *P. berghei* – only infected group. However, the *L. donovani* – only group survived the longest suggesting that *L. donovani* may have conferred some immune protection to the mice in the group. The results were in agreement with other studies by [32, 33] show that mice infected with *P. berghei* only had a high mortality rate as compared to *Leishmania* only infected mice. In addition, it was reported that co-infection decreases the mortality rate during acute *P. yoelii* infection (21).

The findings of the current study demonstrated the presence of functional granulomas in single and co-infected *L. donovani* mice previously done by (31) which serve as a distinctive feature of hepatic resistance. In another study (32) showed that hyaline deposits at times tend to replace disrupted reactive lymphoid follicles when the disease advances. These findings are similar to other works done on characterizing histopathological changes in liver and spleen during canine leishmaniasis (33) in which structural changes such as perisplenitis, accelerated changes in leukocyte numbers and related granulomas were commonly observed in *Leishmania* models (34). Microhaemorrhages observed during cerebral malaria in *P. berghei* only infected mice is presumably due to the rupture of small blood vessels in basal ganglia or subcortical white matter when parasite filled erythrocytes block the blood vessels (35).

Based on the current results, we have demonstrated that it is possible for the two diseases to interact in the same individual. The study results indicate significant differences in body weight, parasitemia, *L. donovani* parasite load and IgG levels. However, Th1 represented by (IFN-γ) and Th2 immune responses represented by IL-4 were not significantly different. Malaria-VL co-infected group showed reduction in disease severity by 21% in malaria and 35.3% in VL during the early phase of the study. Our study therefore concluded that concomitant malaria contributes to disease exacerbation in VL cases as observed in high mortality rate. Nevertheless, the study recommends an integrated approach for malaria and VL screening program where malaria is co-endemic in order to promptly initiate antileishmanial and antimalarial treatments. The insights of the study also emphasize on the need to have a better understanding of both clinical and immunological parameters in co-infected individuals to assist policymakers, clinicians and other stakeholders to develop better strategies for improved disease outcomes.

## Acknowledgments

We thank the Malaria Research and Reference Reagent Resource Center (MR4) for providing us with *Plasmodium berghei* ANKA parasites. The authors would like to acknowledge the staff at Rodent Facility, Animal Science Department at Institute of Primate Research (IPR).

